# Strengthening The Organization and Reporting of Microbiome Studies (STORMS): A Reporting Checklist for Human Microbiome Research

**DOI:** 10.1101/2020.06.24.167353

**Authors:** Chloe Mirzayi, Audrey Renson, Fatima Zohra, Shaimaa Elsafoury, Ludwig Geistlinger, Lora Kasselman, Kelly Eckenrode, Janneke van de Wijgert, Amy Loughman, Francine Z. Marques, STORMS Consortium, Genomic Standards Consortium, Massive Analysis and Quality Control Society, Nicola Segata, Curtis Huttenhower, Jennifer B. Dowd, Heidi E. Jones, Levi Waldron

**Author notes:** Equal contribution (Curtis Huttenhower, Jennifer B. Dowd, Heidi E. Jones, Levi Waldron).

## Abstract

**Background:** Human microbiome research is a growing field with the potential for improving our understanding and treatment of diseases and other conditions. The field is interdisciplinary, making concise organization and reporting of results across different styles of epidemiology, biology, bioinformatics, translational medicine, and statistics a challenge. Commonly used reporting guidelines for observational or genetic epidemiology studies lack key features specific to microbiome studies.

**Methods:** A multidisciplinary group of microbiome epidemiology researchers reviewed elements of available reporting guidelines for observational and genetic studies and adapted these for application to culture-independent human microbiome studies. New reporting elements were developed for laboratory, bioinformatic, and statistical analyses tailored to microbiome studies, and other parts of these checklists were streamlined to keep reporting manageable.

**Results:** STORMS is a 17-item checklist for reporting on human microbiome studies, organized into six sections covering typical sections of a scientific publication, presented as a table with space for author-provided details and intended for inclusion in supplementary materials.

**Conclusions:** STORMS provides guidance for authors and standardization for interdisciplinary microbiome studies, facilitating complete and concise reporting and augments information extraction for downstream applications.

**Availability:** The STORMS checklist is available as a versioned spreadsheet from https://www.stormsmicrobiome.org/.

## Introduction

Changes in the human microbiome have been associated with many disease and health states (1). However, reporting the results of human microbiome research is challenging as it often involves approaches from microbiology, genomics, biomedicine, bioinformatics, statistics, epidemiology, and other fields, resulting in a lack of consistent recommendations for reporting of methods and results. Inconsistent reporting can have consequences for the field by affecting the reproducibility of study results (2). While researchers have called for better reporting standards,(3) such as the Genomic Standards Consortium’s MIxS checklist,(4) to provide a means for reporting sampling, processing and data generation; no comprehensive standardized guidelines spanning laboratory and epidemiological reporting have been proposed.

Standard reporting guidelines promote research consistency and, as a consequence, encourage reproducibility and improved study design. Editorial adoption of the CONsolidated Standards Of Reporting Trials (CONSORT) guidelines, for example, has been associated with an increase in quality of trial reporting.(5,6) Other epidemiological reporting guidelines have seen broad adoption, such as Strengthening the Reporting of OBservational studies in Epidemiology (STROBE)(7) and STrengthening the REporting of Genetic Association Studies (STREGA).(8) STROBE-metagenomics(9) proposes an extension to the STROBE checklist for metagenomics studies. Subsequent to the Minimum Information About a Microarray Experiment (MIAME) (9), Minimum Information about a MARKer gene Sequence (MIMARKS) and Minimum Information about any (x) Sequence (MIxS) checklists provide detailed guidance on reporting of sequencing studies in general. These are focused on the technical aspects of data generation, however, as are projects such as the Microbiome Quality Control Project (MBQC) (10) and International Human Microbiome Standards (IHMS).(11)(12) Together, these serve as useful foundations, but do not span the full range of reporting of human microbiome studies, include items intended for other types of studies, and provide limited guidance on manuscript preparation.

Studies of the human microbiome share many features with other types of molecular epidemiology, but also require unique considerations with their own methodological best practices and reporting standards. In addition to standard elements of epidemiological study design, culture-independent microbiome studies involve collection, handling and preservation of biological specimens, evolving approaches to laboratory processing with elevated potential for batch effects, bioinformatic processing, statistical analysis of sparse, unusually-distributed, high-dimensional data, and reporting of results on potentially thousands of microbial features.(13–15) Because there is no agreed-upon gold-standard method for microbiome research and the field has not reached consensus on many of these aspects, inconsistencies in reporting inhibit reproducibility and hamper efforts to draw conclusions across similar studies.

For these reasons, we convened a multi-disciplinary working group to develop guidelines tailored to microbiome study reporting. Members of this group include epidemiologists, biostatisticians, bioinformaticians, physician-scientists, genomicists, and microbiologists. The checklist is designed to balance completeness with burden of use, and is applicable to a broad range of human microbiome study designs and analysis. The “Strengthening The Organization and Reporting of Microbiome Studies (STORMS)” checklist draws relevant items from related guidelines and adds new tailored guidelines to serve as a tool to organize study planning and manuscript preparation, to improve the clarity of manuscripts, and to facilitate reviewers and readers in assessing these studies.

## Methods

### Origin and development

The origins of these guidelines are rooted in a project to create a standardized database of published literature reporting relationships between the microbiome and disease. The goal of that project is to create a publicly available, standardized database of microbiome study findings indexed by condition of interest (e.g. disease, health status, diet, environmental factor) microbiome site (e.g. gut, mouth, skin), and microbial taxonomy to aid comparative analysis. As of August 2020, 22 curators have extracted findings from 371 unique published studies. Included studies must have examined the relationship between the microbiome and a condition of interest and included findings on a taxonomic level (even if all findings were null).

This review revealed substantial reporting heterogeneity, particularly for epidemiology, such as study design, confounding factors, and sources of bias. It also revealed microbiome-specific issues, including statistical analysis of compositional relative abundance data and handling ‘batch’ effects.(16) This heterogeneity highlighted the need for standardized reporting guidelines, similar to those used in other fields of study. The curators determined that standardized reporting guidelines would streamline the review process, but would more importantly help researchers throughout the field of microbiome research communicate their findings effectively.

The resulting multidisciplinary group of bioinformaticians, epidemiologists, biostatisticians, and microbiologists was thus convened to discuss microbiome reporting standards. The group began by reviewing existing reporting standards including STROBE,(17) STROBE-ME,(18) STREGA,(8) MICRO,(19), MIMARKS,(4) and STROGAR.(20) The group also reviewed existing articles containing recommendations for microbiome reporting.(21,22) The STROBE and STREGA guidelines were used as a starting point for the STORMS checklist, although aspects were incorporated from the other reporting standards.

Following the reporting standards development guidelines recommended by EQUATOR, a comprehensive list of potential guideline items was created. From this list, group members added, modified, and removed items based on their expertise. After the first round of edits, the checklist was then applied to a recent microbiome study(23) by group members. Comments, removals, and additions were harmonized after each round. Based on this process, additional changes, simplifications, and clarifications were made. This process was repeated until there was a group consensus that the checklist was ready for use.

In addition to the core working group, outside subject matter experts identified by working group members were then invited to review the guidelines and provide feedback as members of the STORMS Consortium. In total, 113 experts were invited to join the STORMS Consortium and 61 accepted and provided feedback. Substantive feedback (i.e. not grammar, spelling, or other small changes) was compiled into a feedback document (Supplement 1) and the core working group responded to each piece of feedback individually. Additional revisions were made based on their comments. After this round of revisions, consortium members were once again invited to review the checklist prior to submission for publication.

## Results

### Checklist

The latest version of the checklist at time of publication is presented in Table 1 and a summary of items is presented in Figure 1. Of the items in the latest version in the STORMS checklist, nine items or sub-items were unchanged from STROBE, three were modified from STROBE, one was modified from STREGA, and 57 new guidelines were developed. Nine Items with an overlap with MIxS are specified. Rationale for new and modified items are presented below. Documentation of items unmodified from STROBE and STREGA were presented in the publications of those checklists.

**Table 1.**
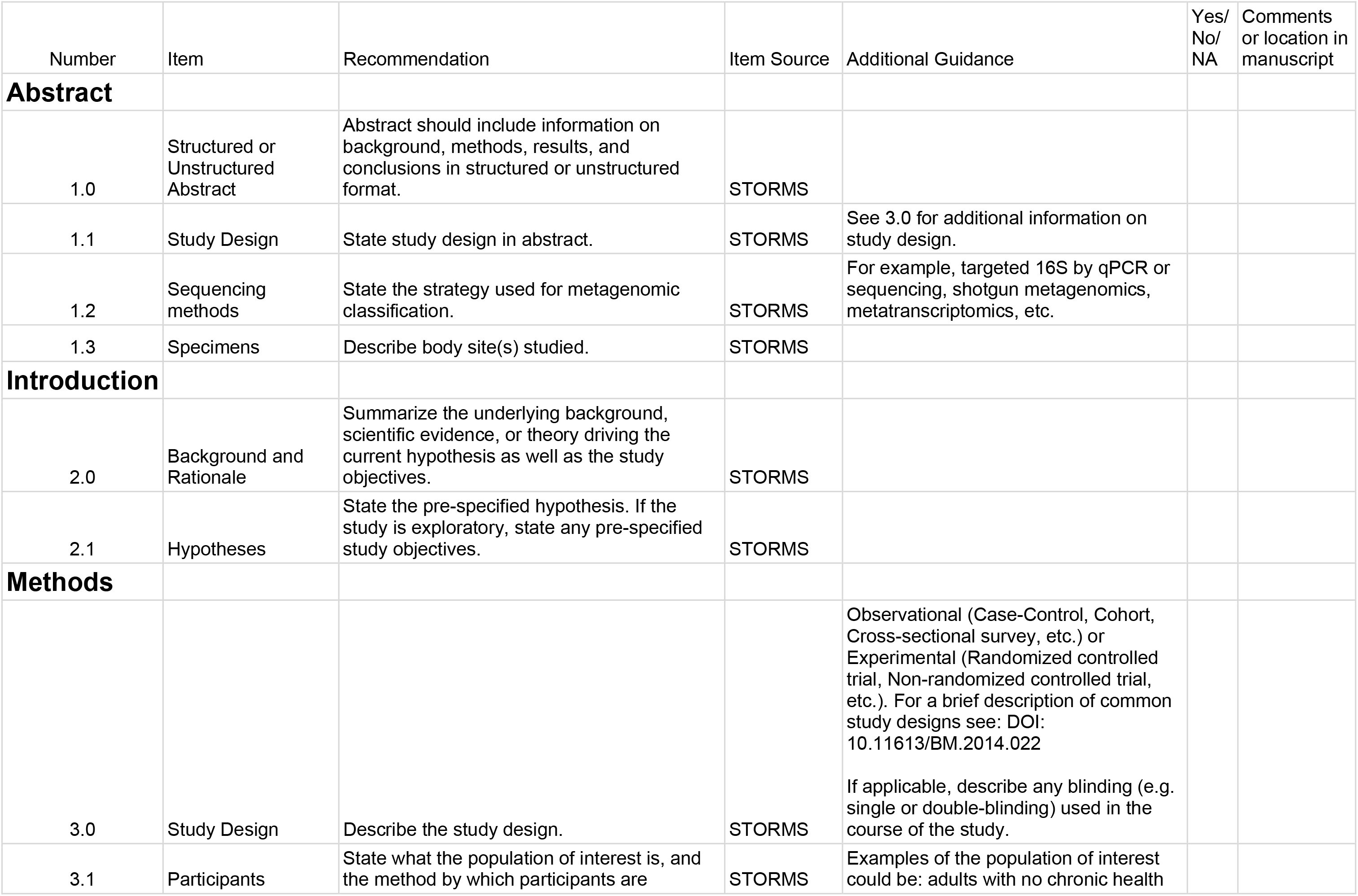
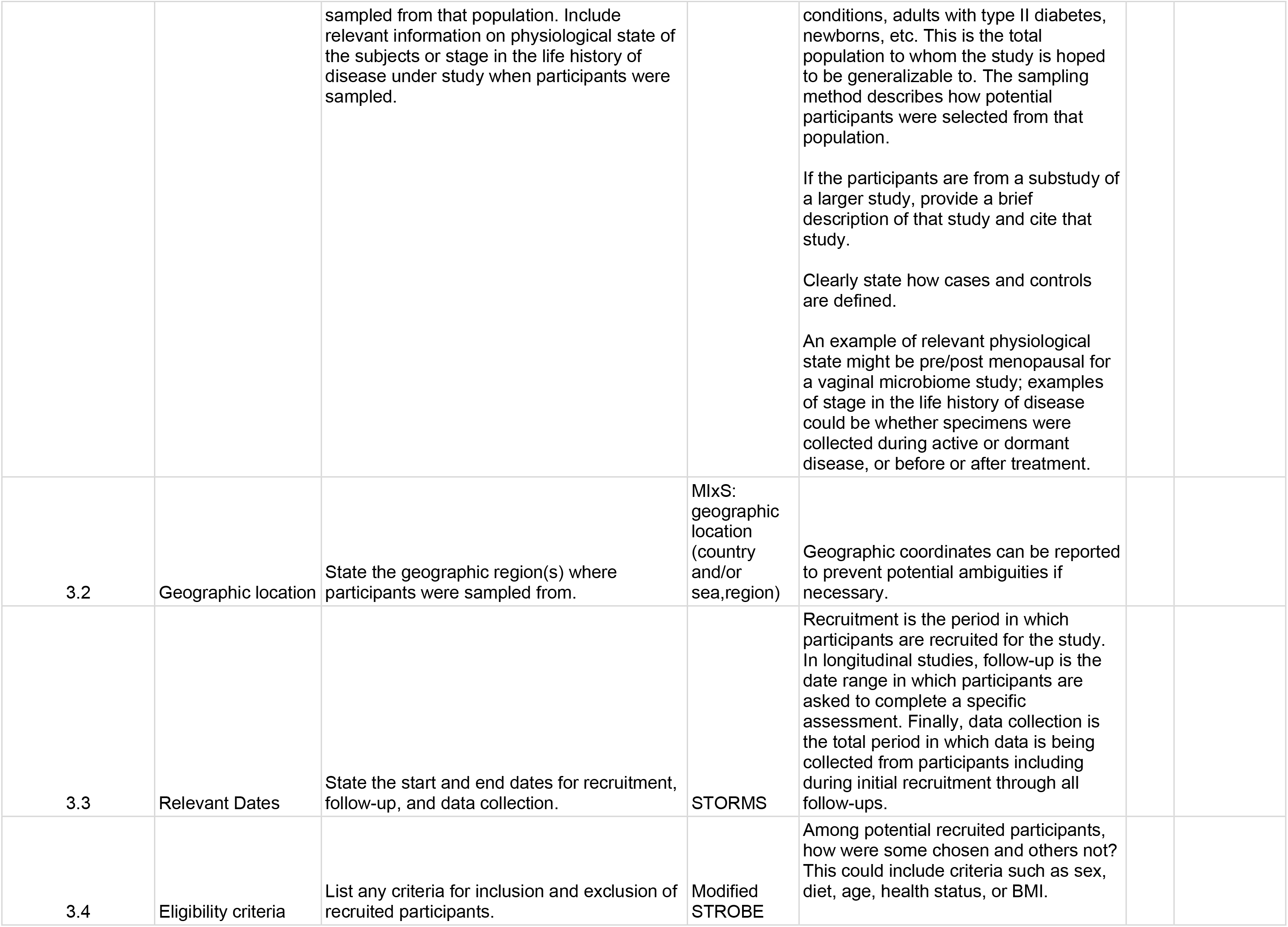
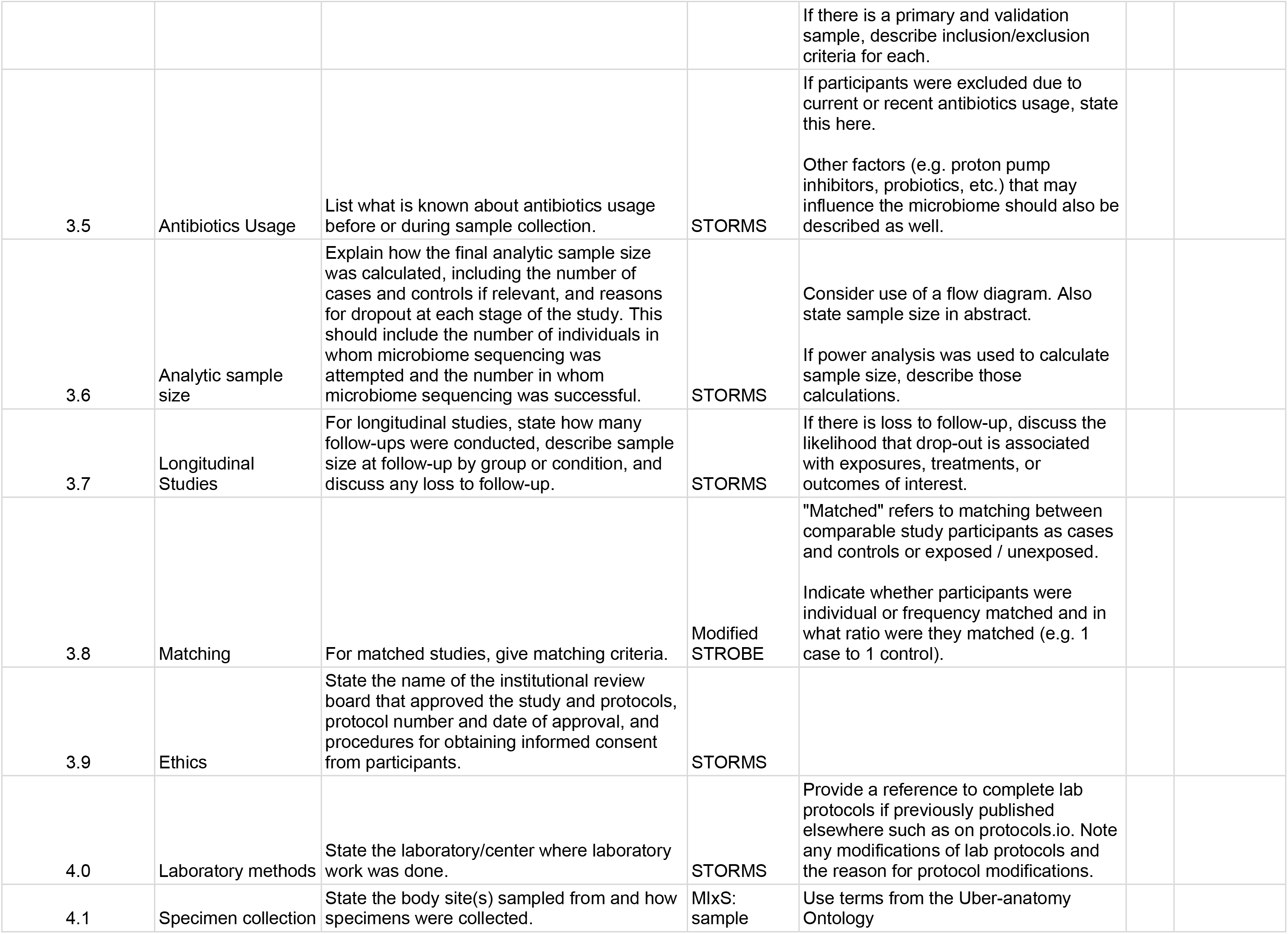
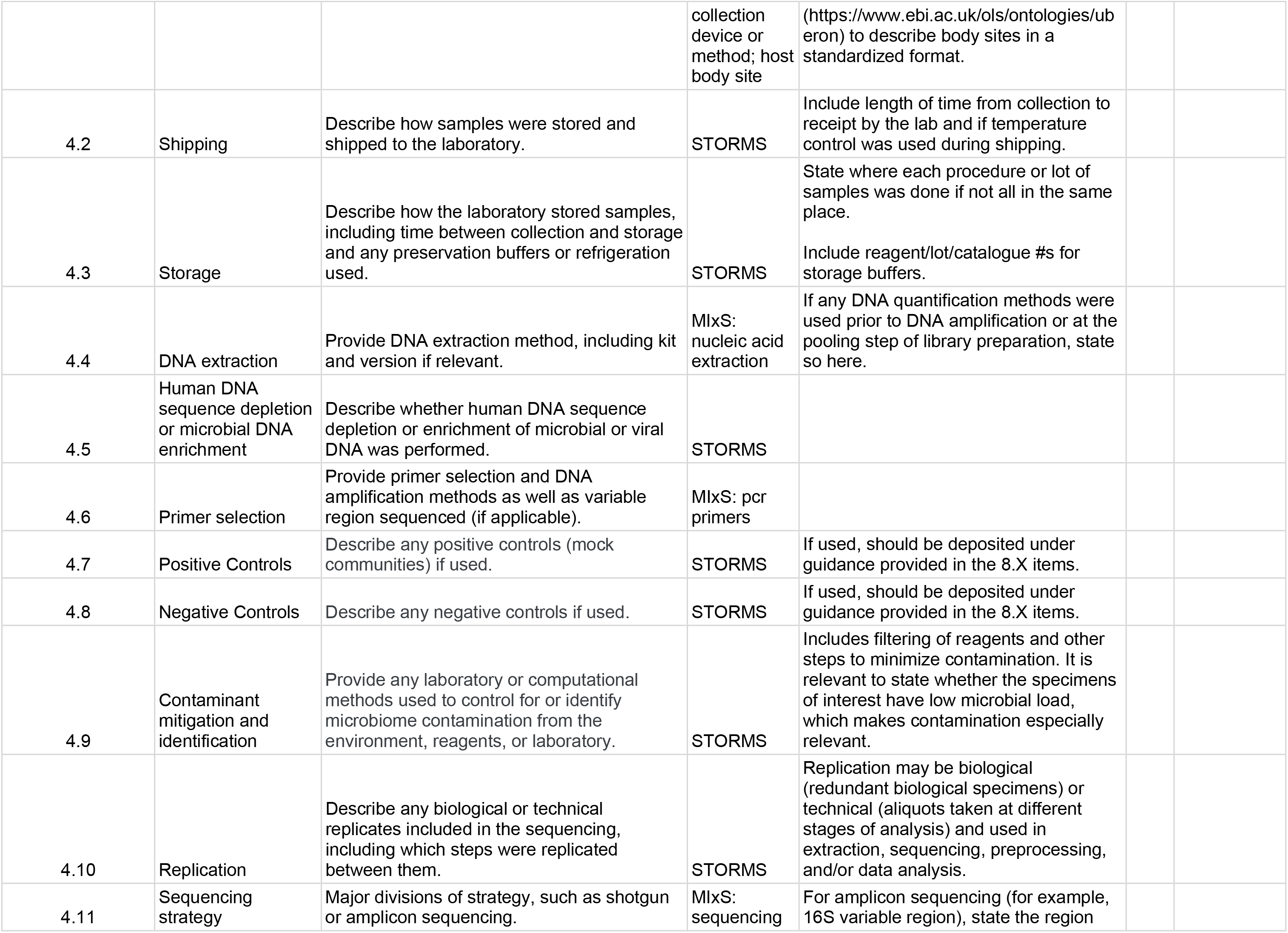
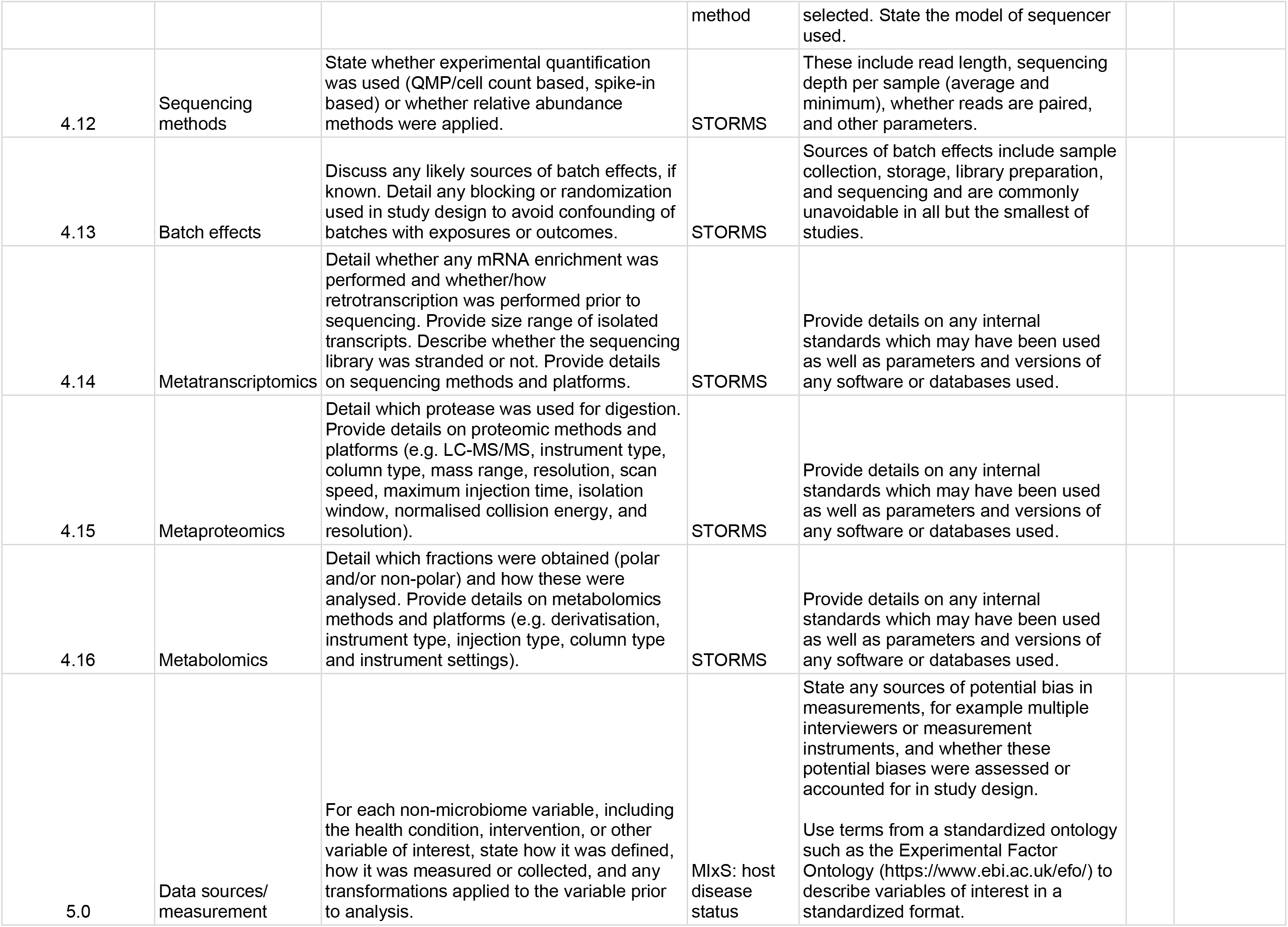
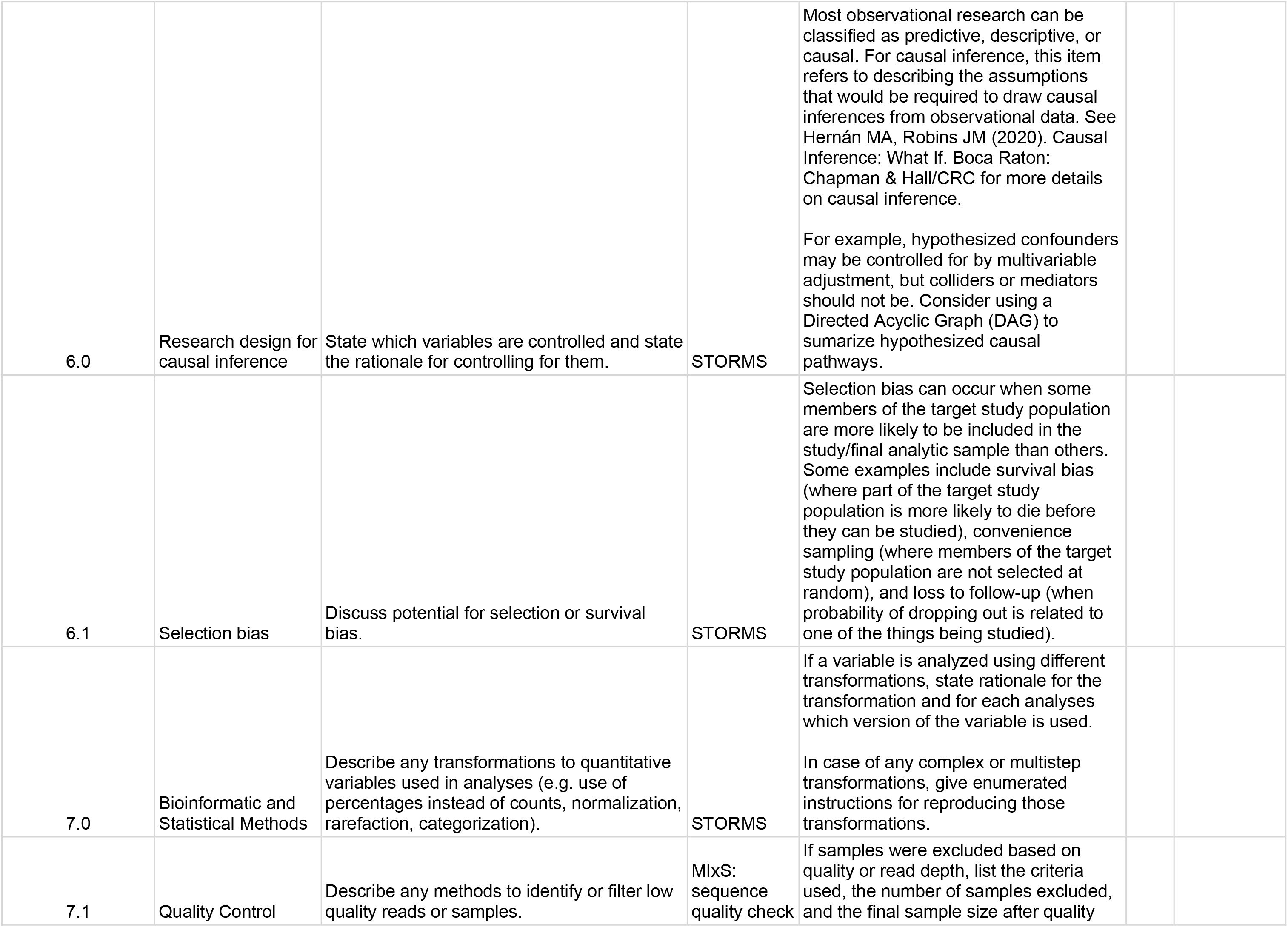
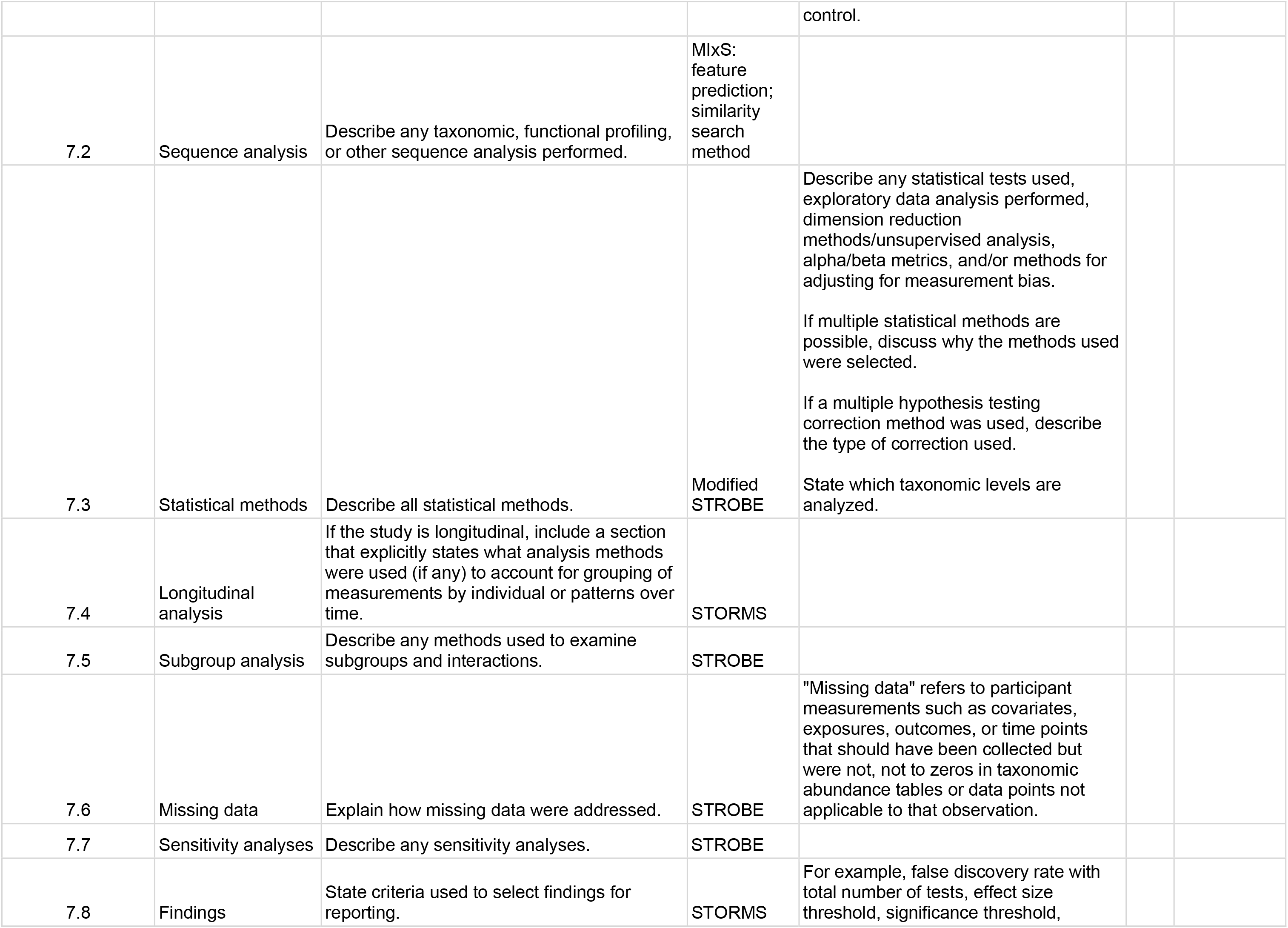
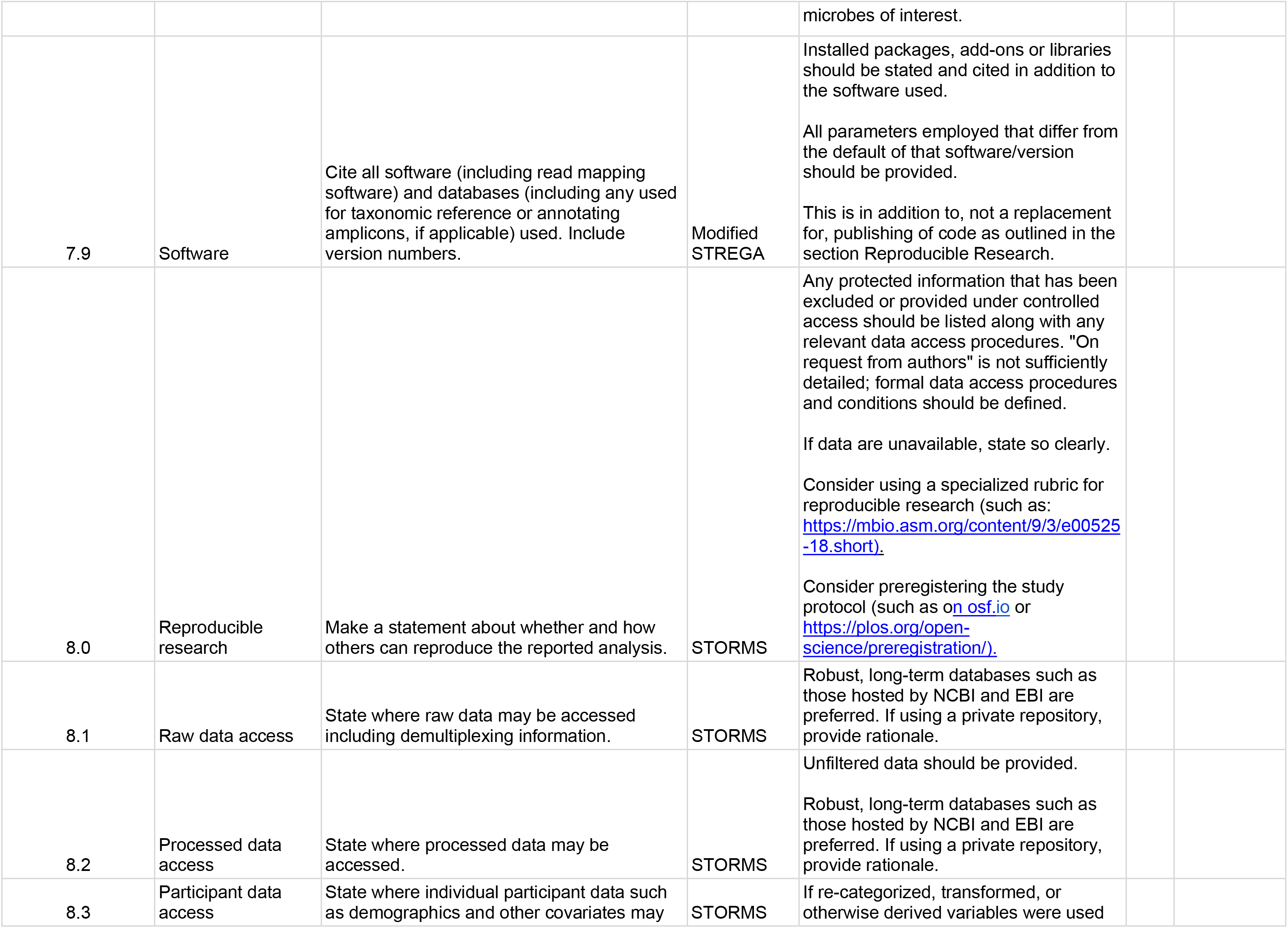
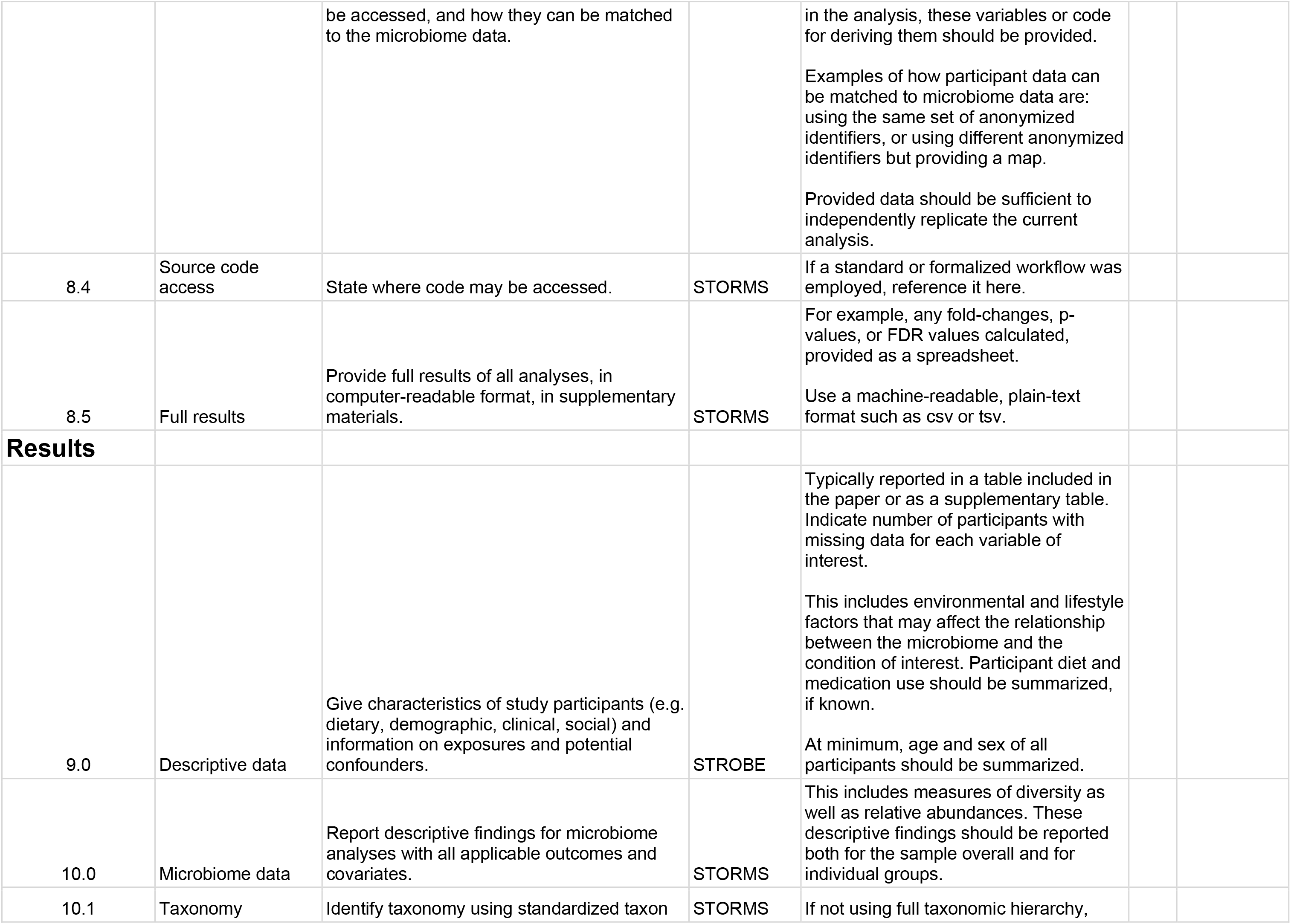
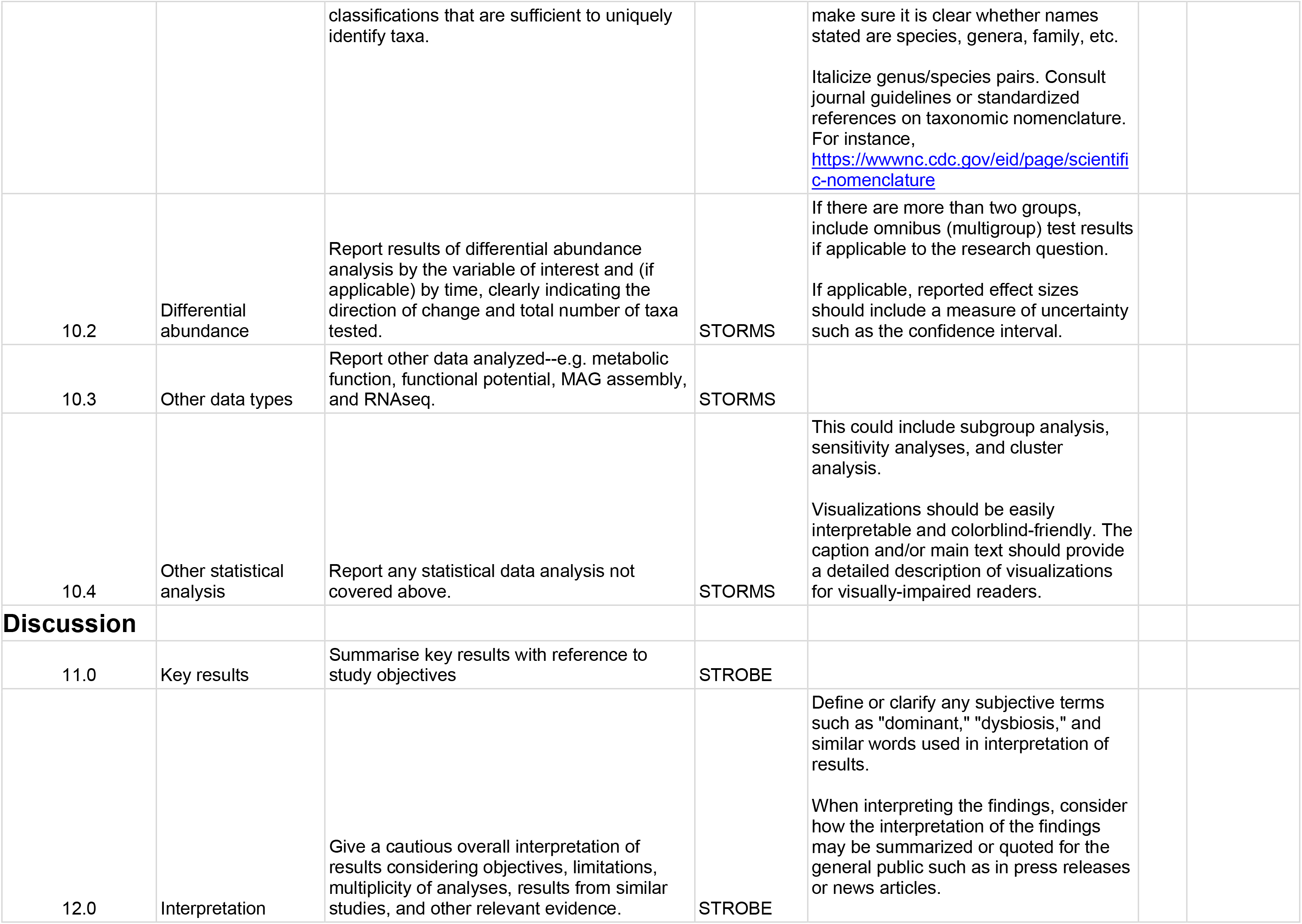
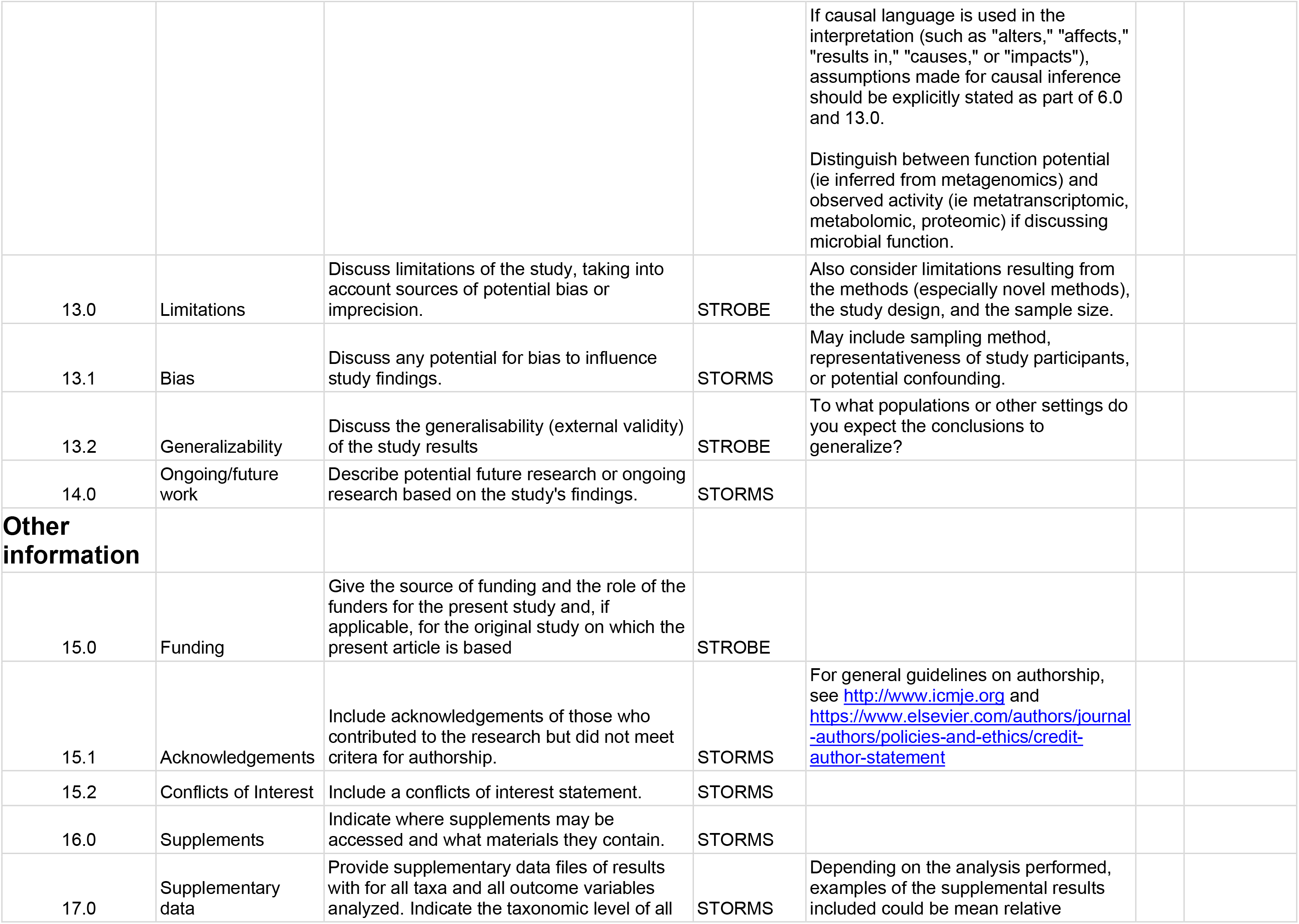
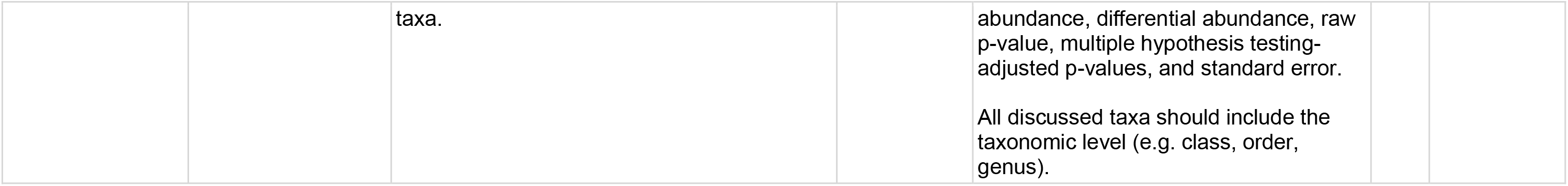
STORMS Checklist. For the latest version of the checklist, please visit https://www.stormsmicrobiome.org/.

| Number | Item | Recommendation | Item Source | Additional Guidance | Yes/<br>No/<br>NA | Comments<br>or location in<br>manuscript |
| --- | --- | --- | --- | --- | --- | --- |
| **Abstract** |  |  |  |  |  |  |
| 1.0 | Structured or Unstructured Abstract | Abstract should include information on background, methods, results, and conclusions in structured or unstructured format. | STORMS |  |  |  |
| 1.1 | Study Design | State study design in abstract. | STORMS | See 3.0 for additional information on study design. |  |  |
| 1.2 | Sequencing methods | State the strategy used for metagenomic classification. | STORMS | For example, targeted 16S by qPCR or sequencing, shotgun metagenomics, metatranscriptomics, etc. |  |  |
| 1.3 | Specimens | Describe body site(s) studied. | STORMS |  |  |  |
| **Introduction** |  |  |  |  |  |  |
| 2.0 | Background and Rationale | Summarize the underlying background, scientific evidence, or theory driving the current hypothesis as well as the study objectives. | STORMS |  |  |  |
| 2.1 | Hypotheses | State the pre-specified hypothesis. If the study is exploratory, state any pre-specified study objectives. | STORMS |  |  |  |
| **Methods** |  |  |  |  |  |  |
| 3.0 | Study Design | Describe the study design. | STORMS | Observational (Case-Control, Cohort, Cross-sectional survey, etc.) or Experimental (Randomized controlled trial, Non-randomized controlled trial, etc.). For a brief description of common study designs see: DOI: 10.11613/BM.2014.022<br>If applicable, describe any blinding (e.g. single or double-blinding) used in the course of the study. |  |  |
| 3.1 | Participants | State what the population of interest is, and the method by which participants are | STORMS | Examples of the population of interest could be: adults with no chronic health |  |  |
|  |  | sampled from that population. Include relevant information on physiological state of the subjects or stage in the life history of disease under study when participants were sampled. |  | conditions, adults with type II diabetes, newborns, etc. This is the total population to whom the study is hoped to be generalizable to. The sampling method describes how potential participants were selected from that population.<br>If the participants are from a substudy of a larger study, provide a brief description of that study and cite that study.<br>Clearly state how cases and controls are defined.<br>An example of relevant physiological state might be pre/post menopausal for a vaginal microbiome study; examples of stage in the life history of disease could be whether specimens were collected during active or dormant disease, or before or after treatment. |  |  |
| 3.2 | Geographic location | State the geographic region(s) where participants were sampled from. | MIxS:<br>geographic location (country and/or sea,region) | Geographic coordinates can be reported to prevent potential ambiguities if necessary. |  |  |
| 3.3 | Relevant Dates | State the start and end dates for recruitment, follow-up, and data collection. | STORMS | Recruitment is the period in which participants are recruited for the study. In longitudinal studies, follow-up is the date range in which participants are asked to complete a specific assessment. Finally, data collection is the total period in which data is being collected from participants including during initial recruitment through all follow-ups. |  |  |
| 3.4 | Eligibility criteria | List any criteria for inclusion and exclusion of recruited participants. | Modified STROBE | Among potential recruited participants, how were some chosen and others not? This could include criteria such as sex, diet, age, health status, or BMI. |  |  |
|  |  |  |  | If there is a primary and validation sample, describe inclusion/exclusion criteria for each. |  |  |
| 3.5 | Antibiotics Usage | List what is known about antibiotics usage before or during sample collection. | STORMS | If participants were excluded due to current or recent antibiotics usage, state this here.<br>Other factors (e.g. proton pump inhibitors, probiotics, etc.) that may influence the microbiome should also be described as well. |  |  |
| 3.6 | Analytic sample size | Explain how the final analytic sample size was calculated, including the number of cases and controls if relevant, and reasons for dropout at each stage of the study. This should include the number of individuals in whom microbiome sequencing was attempted and the number in whom microbiome sequencing was successful. | STORMS | Consider use of a flow diagram. Also state sample size in abstract.<br>If power analysis was used to calculate sample size, describe those calculations. |  |  |
| 3.7 | Longitudinal Studies | For longitudinal studies, state how many follow-ups were conducted, describe sample size at follow-up by group or condition, and discuss any loss to follow-up. | STORMS | If there is loss to follow-up, discuss the likelihood that drop-out is associated with exposures, treatments, or outcomes of interest. |  |  |
| 3.8 | Matching | For matched studies, give matching criteria. | Modified STROBE | "Matched" refers to matching between comparable study participants as cases and controls or exposed / unexposed.<br>Indicate whether participants were individual or frequency matched and in what ratio were they matched (e.g. 1 case to 1 control). |  |  |
| 3.9 | Ethics | State the name of the institutional review board that approved the study and protocols, protocol number and date of approval, and procedures for obtaining informed consent from participants. | STORMS |  |  |  |
| 4.0 | Laboratory methods | State the laboratory/center where laboratory work was done. | STORMS | Provide a reference to complete lab protocols if previously published elsewhere such as on protocols.io. Note any modifications of lab protocols and the reason for protocol modifications. |  |  |
| 4.1 | Specimen collection | State the body site(s) sampled from and how specimens were collected. | MixS: sample | Use terms from the Uber-anatomy Ontology |  |  |
|  |  |  | collection device or method; host body site | (https://www.ebi.ac.uk/ols/ontologies/uberon) to describe body sites in a standardized format. |  |  |
| 4.2 | Shipping | Describe how samples were stored and shipped to the laboratory. | STORMS | Include length of time from collection to receipt by the lab and if temperature control was used during shipping. |  |  |
| 4.3 | Storage | Describe how the laboratory stored samples, including time between collection and storage and any preservation buffers or refrigeration used. | STORMS | State where each procedure or lot of samples was done if not all in the same place.<br>Include reagent/lot/catalogue #s for storage buffers. |  |  |
| 4.4 | DNA extraction | Provide DNA extraction method, including kit and version if relevant. | MIxS: nucleic acid extraction | If any DNA quantification methods were used prior to DNA amplification or at the pooling step of library preparation, state so here. |  |  |
| 4.5 | Human DNA sequence depletion or microbial DNA enrichment | Describe whether human DNA sequence depletion or enrichment of microbial or viral DNA was performed. | STORMS |  |  |  |
| 4.6 | Primer selection | Provide primer selection and DNA amplification methods as well as variable region sequenced (if applicable). | MIxS: pcr primers |  |  |  |
| 4.7 | Positive Controls | Describe any positive controls (mock communities) if used. | STORMS | If used, should be deposited under guidance provided in the 8.X items. |  |  |
| 4.8 | Negative Controls | Describe any negative controls if used. | STORMS | If used, should be deposited under guidance provided in the 8.X items. |  |  |
| 4.9 | Contaminant mitigation and identification | Provide any laboratory or computational methods used to control for or identify microbiome contamination from the environment, reagents, or laboratory. | STORMS | Includes filtering of reagents and other steps to minimize contamination. It is relevant to state whether the specimens of interest have low microbial load, which makes contamination especially relevant. |  |  |
| 4.10 | Replication | Describe any biological or technical replicates included in the sequencing, including which steps were replicated between them. | STORMS | Replication may be biological (redundant biological specimens) or technical (aliquots taken at different stages of analysis) and used in extraction, sequencing, preprocessing, and/or data analysis. |  |  |
| 4.11 | Sequencing strategy | Major divisions of strategy, such as shotgun or amplicon sequencing. | MIxS: sequencing | For amplicon sequencing (for example, 16S variable region), state the region |  |  |
|  |  |  | method | selected. State the model of sequencer used. |  |  |
| 4.12 | Sequencing methods | State whether experimental quantification was used (QMP/cell count based, spike-in based) or whether relative abundance methods were applied. | STORMS | These include read length, sequencing depth per sample (average and minimum), whether reads are paired, and other parameters. |  |  |
| 4.13 | Batch effects | Discuss any likely sources of batch effects, if known. Detail any blocking or randomization used in study design to avoid confounding of batches with exposures or outcomes. | STORMS | Sources of batch effects include sample collection, storage, library preparation, and sequencing and are commonly unavoidable in all but the smallest of studies. |  |  |
| 4.14 | Metatranscriptomics | Detail whether any mRNA enrichment was performed and whether/how retrotranscription was performed prior to sequencing. Provide size range of isolated transcripts. Describe whether the sequencing library was stranded or not. Provide details on sequencing methods and platforms. | STORMS | Provide details on any internal standards which may have been used as well as parameters and versions of any software or databases used. |  |  |
| 4.15 | Metaproteomics | Detail which protease was used for digestion. Provide details on proteomic methods and platforms (e.g. LC-MS/MS, instrument type, column type, mass range, resolution, scan speed, maximum injection time, isolation window, normalised collision energy, and resolution). | STORMS | Provide details on any internal standards which may have been used as well as parameters and versions of any software or databases used. |  |  |
| 4.16 | Metabolomics | Detail which fractions were obtained (polar and/or non-polar) and how these were analysed. Provide details on metabolomics methods and platforms (e.g. derivatisation, instrument type, injection type, column type and instrument settings). | STORMS | Provide details on any internal standards which may have been used as well as parameters and versions of any software or databases used. |  |  |
| 5.0 | Data sources/ measurement | For each non-microbiome variable, including the health condition, intervention, or other variable of interest, state how it was defined, how it was measured or collected, and any transformations applied to the variable prior to analysis. | MIxS: host disease status | State any sources of potential bias in measurements, for example multiple interviewers or measurement instruments, and whether these potential biases were assessed or accounted for in study design.<br>Use terms from a standardized ontology such as the Experimental Factor Ontology (https://www.ebi.ac.uk/efo/) to describe variables of interest in a standardized format. |  |  |
| 6.0 | Research design for causal inference | State which variables are controlled and state the rationale for controlling for them. | STORMS | Most observational research can be classified as predictive, descriptive, or causal. For causal inference, this item refers to describing the assumptions that would be required to draw causal inferences from observational data. See Hernán MA, Robins JM (2020). Causal Inference: What If. Boca Raton: Chapman & Hall/CRC for more details on causal inference.<br>For example, hypothesized confounders may be controlled for by multivariable adjustment, but colliders or mediators should not be. Consider using a Directed Acyclic Graph (DAG) to summarize hypothesized causal pathways. |  |  |
| 6.1 | Selection bias | Discuss potential for selection or survival bias. | STORMS | Selection bias can occur when some members of the target study population are more likely to be included in the study/final analytic sample than others. Some examples include survival bias (where part of the target study population is more likely to die before they can be studied), convenience sampling (where members of the target study population are not selected at random), and loss to follow-up (when probability of dropping out is related to one of the things being studied). |  |  |
| 7.0 | Bioinformatic and Statistical Methods | Describe any transformations to quantitative variables used in analyses (e.g. use of percentages instead of counts, normalization, rarefaction, categorization). | STORMS | If a variable is analyzed using different transformations, state rationale for the transformation and for each analyses which version of the variable is used.<br>In case of any complex or multistep transformations, give enumerated instructions for reproducing those transformations. |  |  |
| 7.1 | Quality Control | Describe any methods to identify or filter low quality reads or samples. | MIxS: sequence quality check | If samples were excluded based on quality or read depth, list the criteria used, the number of samples excluded, and the final sample size after quality control. |  |  |
| 7.2 | Sequence analysis | Describe any taxonomic, functional profiling, or other sequence analysis performed. | MixS:<br>feature<br>prediction;<br>similarity<br>search<br>method |  |  |  |
| 7.3 | Statistical methods | Describe all statistical methods. | Modified<br>STROBE | Describe any statistical tests used, exploratory data analysis performed, dimension reduction methods/unsupervised analysis, alpha/beta metrics, and/or methods for adjusting for measurement bias.<br>If multiple statistical methods are possible, discuss why the methods used were selected.<br>If a multiple hypothesis testing correction method was used, describe the type of correction used.<br>State which taxonomic levels are analyzed. |  |  |
| 7.4 | Longitudinal analysis | If the study is longitudinal, include a section that explicitly states what analysis methods were used (if any) to account for grouping of measurements by individual or patterns over time. | STORMS |  |  |  |
| 7.5 | Subgroup analysis | Describe any methods used to examine subgroups and interactions. | STROBE |  |  |  |
| 7.6 | Missing data | Explain how missing data were addressed. | STROBE | "Missing data" refers to participant measurements such as covariates, exposures, outcomes, or time points that should have been collected but were not, not to zeros in taxonomic abundance tables or data points not applicable to that observation. |  |  |
| 7.7 | Sensitivity analyses | Describe any sensitivity analyses. | STROBE |  |  |  |
| 7.8 | Findings | State criteria used to select findings for reporting. | STORMS | For example, false discovery rate with total number of tests, effect size threshold, significance threshold, microbes of interest. |  |  |
| 7.9 | Software | Cite all software (including read mapping software) and databases (including any used for taxonomic reference or annotating amplicons, if applicable) used. Include version numbers. | Modified STREGA | Installed packages, add-ons or libraries should be stated and cited in addition to the software used.<br>All parameters employed that differ from the default of that software/version should be provided.<br>This is in addition to, not a replacement for, publishing of code as outlined in the section Reproducible Research. |  |  |
| 8.0 | Reproducible research | Make a statement about whether and how others can reproduce the reported analysis. | STORMS | Any protected information that has been excluded or provided under controlled access should be listed along with any relevant data access procedures. "On request from authors" is not sufficiently detailed; formal data access procedures and conditions should be defined.<br>If data are unavailable, state so clearly.<br>Consider using a specialized rubric for reproducible research (such as: https://mbio.asm.org/content/9/3/e00525-18.short).<br>Consider preregistering the study protocol (such as on osf.io or https://plos.org/open-science/preregistration/). |  |  |
| 8.1 | Raw data access | State where raw data may be accessed including demultiplexing information. | STORMS | Robust, long-term databases such as those hosted by NCBI and EBI are preferred. If using a private repository, provide rationale. |  |  |
| 8.2 | Processed data access | State where processed data may be accessed. | STORMS | Unfiltered data should be provided.<br>Robust, long-term databases such as those hosted by NCBI and EBI are preferred. If using a private repository, provide rationale. |  |  |
| 8.3 | Participant data access | State where individual participant data such as demographics and other covariates may | STORMS | If re-categorized, transformed, or otherwise derived variables were used |  |  |
|  |  | be accessed, and how they can be matched to the microbiome data. |  | in the analysis, these variables or code for deriving them should be provided.<br>Examples of how participant data can be matched to microbiome data are: using the same set of anonymized identifiers, or using different anonymized identifiers but providing a map.<br>Provided data should be sufficient to independently replicate the current analysis. |  |  |
| 8.4 | Source code access | State where code may be accessed. | STORMS | If a standard or formalized workflow was employed, reference it here. |  |  |
| 8.5 | Full results | Provide full results of all analyses, in computer-readable format, in supplementary materials. | STORMS | For example, any fold-changes, p-values, or FDR values calculated, provided as a spreadsheet.<br>Use a machine-readable, plain-text format such as csv or tsv. |  |  |
| **Results** |  |  |  |  |  |  |
| 9.0 | Descriptive data | Give characteristics of study participants (e.g. dietary, demographic, clinical, social) and information on exposures and potential confounders. | STROBE | Typically reported in a table included in the paper or as a supplementary table. Indicate number of participants with missing data for each variable of interest.<br>This includes environmental and lifestyle factors that may affect the relationship between the microbiome and the condition of interest. Participant diet and medication use should be summarized, if known.<br>At minimum, age and sex of all participants should be summarized. |  |  |
| 10.0 | Microbiome data | Report descriptive findings for microbiome analyses with all applicable outcomes and covariates. | STORMS | This includes measures of diversity as well as relative abundances. These descriptive findings should be reported both for the sample overall and for individual groups. |  |  |
| 10.1 | Taxonomy | Identify taxonomy using standardized taxon | STORMS | If not using full taxonomic hierarchy, |  |  |
|  |  | classifications that are sufficient to uniquely identify taxa. |  | make sure it is clear whether names stated are species, genera, family, etc.<br>Italicize genus/species pairs. Consult journal guidelines or standardized references on taxonomic nomenclature. For instance, https://wwwnc.cdc.gov/eid/page/scientific-nomenclature |  |  |
| 10.2 | Differential abundance | Report results of differential abundance analysis by the variable of interest and (if applicable) by time, clearly indicating the direction of change and total number of taxa tested. | STORMS | If there are more than two groups, include omnibus (multigroup) test results if applicable to the research question.<br>If applicable, reported effect sizes should include a measure of uncertainty such as the confidence interval. |  |  |
| 10.3 | Other data types | Report other data analyzed--e.g. metabolic function, functional potential, MAG assembly, and RNAseq. | STORMS |  |  |  |
| 10.4 | Other statistical analysis | Report any statistical data analysis not covered above. | STORMS | This could include subgroup analysis, sensitivity analyses, and cluster analysis.<br>Visualizations should be easily interpretable and colorblind-friendly. The caption and/or main text should provide a detailed description of visualizations for visually-impaired readers. |  |  |
| **Discussion** |  |  |  |  |  |  |
| 11.0 | Key results | Summarise key results with reference to study objectives | STROBE |  |  |  |
| 12.0 | Interpretation | Give a cautious overall interpretation of results considering objectives, limitations, multiplicity of analyses, results from similar studies, and other relevant evidence. | STROBE | Define or clarify any subjective terms such as "dominant," "dysbiosis," and similar words used in interpretation of results.<br>When interpreting the findings, consider how the interpretation of the findings may be summarized or quoted for the general public such as in press releases or news articles. |  |  |
|  |  |  |  | If causal language is used in the interpretation (such as "alters," "affects," "results in," "causes," or "impacts"), assumptions made for causal inference should be explicitly stated as part of 6.0 and 13.0.<br>Distinguish between function potential (ie inferred from metagenomics) and observed activity (ie metatranscriptomic, metabolomic, proteomic) if discussing microbial function. |  |  |
| 13.0 | Limitations | Discuss limitations of the study, taking into account sources of potential bias or imprecision. | STROBE | Also consider limitations resulting from the methods (especially novel methods), the study design, and the sample size. |  |  |
| 13.1 | Bias | Discuss any potential for bias to influence study findings. | STORMS | May include sampling method, representativeness of study participants, or potential confounding. |  |  |
| 13.2 | Generalizability | Discuss the generalisability (external validity) of the study results | STROBE | To what populations or other settings do you expect the conclusions to generalize? |  |  |
| 14.0 | Ongoing/future work | Describe potential future research or ongoing research based on the study's findings. | STORMS |  |  |  |
| **Other information** |  |  |  |  |  |  |
| 15.0 | Funding | Give the source of funding and the role of the funders for the present study and, if applicable, for the original study on which the present article is based | STROBE |  |  |  |
| 15.1 | Acknowledgements | Include acknowledgements of those who contributed to the research but did not meet criteria for authorship. | STORMS | For general guidelines on authorship, see http://www.icmje.org and https://www.elsevier.com/authors/journal-authors/policies-and-ethics/credit-author-statement |  |  |
| 15.2 | Conflicts of Interest | Include a conflicts of interest statement. | STORMS |  |  |  |
| 16.0 | Supplements | Indicate where supplements may be accessed and what materials they contain. | STORMS |  |  |  |
| 17.0 | Supplementary data | Provide supplementary data files of results with for all taxa and all outcome variables analyzed. Indicate the taxonomic level of all | STORMS | Depending on the analysis performed, examples of the supplemental results included could be mean relative |  |  |
|  |  | taxa. |  | abundance, differential abundance, raw p-value, multiple hypothesis testing-adjusted p-values, and standard error.<br>All discussed taxa should include the taxonomic level (e.g. class, order, genus). |  |  |

**Figure 1.**
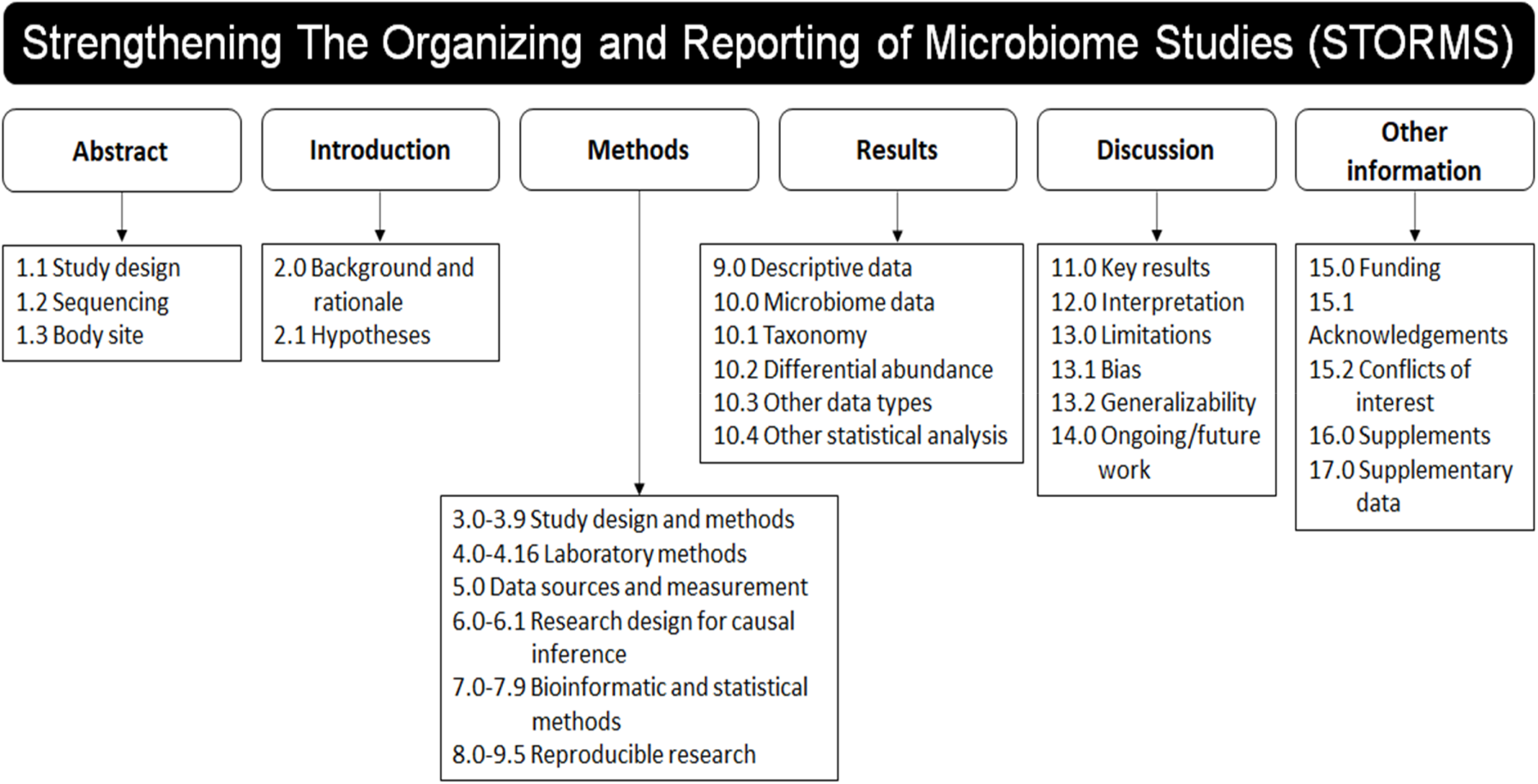
STORMS checklist items by major manuscript heading. For detailed descriptions of each item and additional guidance, see the STORMS checklist (Table 1).

#### Abstract (1.0-1.3)

Along with commonly included abstract materials such as a basic description of the participants and results, authors should report the study design(24)--such as cross-sectional, case-control, cohort, or randomized controlled trial--in the abstract of their article (item 1.1), as required by other reporting guidelines. Communicating study design in the abstract allows readers to quickly categorize the type of evidence provided. As part of this basic description, sequencing methods should be mentioned (item 1.2). Body site(s) sampled should also be included (item 1.3).

#### Introduction (2.0-2.1)

The introduction should clearly describe the underlying background, evidence, or theory that motivated the current study (item 2.0). Among other possibilities, this could include pilot study data, previous findings from a similar study or topic, or a biologically plausible mechanism that has been proposed. This clarifies for the reader the motivations for the present study. If the study is exploratory in nature, it should be explained what motivated the current exploration and the goals of the exploratory study. The hypothesis developed based on the background should be included. If the study was exploratory and did not define a hypothesis, pre-specified study objectives should be included (item 2.1).

#### Methods (3.0-8.5)

##### Participants (3.0-3.9)

The methods section should contain sufficient information for study replicability. Because study design is essential to understanding the study, it should be stated in the methods (item 3.0). When describing the participants in the study, the population of interest should be described and how participants were sampled from the source population (item 3.1). Because participant characteristics such as environment,(25) lifestyle behaviors, diet, biomedical interventions, demographics,(26) and geography(27) (item 3.2) can correspond with substantial differences in the microbiome, it is essential to include this description. Temporal context can be quite important as well so start and end dates for recruitment, follow-up, and data collection should be stated (item 3.3).

Specific criteria used to assess potential participants for eligibility in the study should also be reported, detailing both inclusion and exclusion criteria (item 3.4). Inclusion and exclusion criteria are pre-established characteristics used for selection of participants into the study, and describing these criteria is essential in understanding the study’s target population.(28) This is expanded from STROBE, which requires eligibility criteria, but does not specify that both inclusion and exclusion criteria should be reported in detail. Any information collected about antibiotics or other treatments that could affect the microbiome should be described (item 3.5) as well as if any exclusion criteria included recent antibiotic or other medication use.

The final analytic sample sizes and read numbers should be stated as well as the reason for any exclusion of participants at any step of the recruitment, follow-up, or laboratory processes (item 3.6). STROBE suggests using a flow diagram to show when and why participants were removed from the study. Examples of such flow diagrams are presented in Figure 2. If participants were lost to follow-up or did not complete all assessments in a longitudinal study, details on how follow-ups were conducted should be stated and time-point specific sample sizes should also be reported (item 3.7). Additionally, studies that matched cases to controls should describe what variables were used in matching (item 3.4).

**Figure 2.**
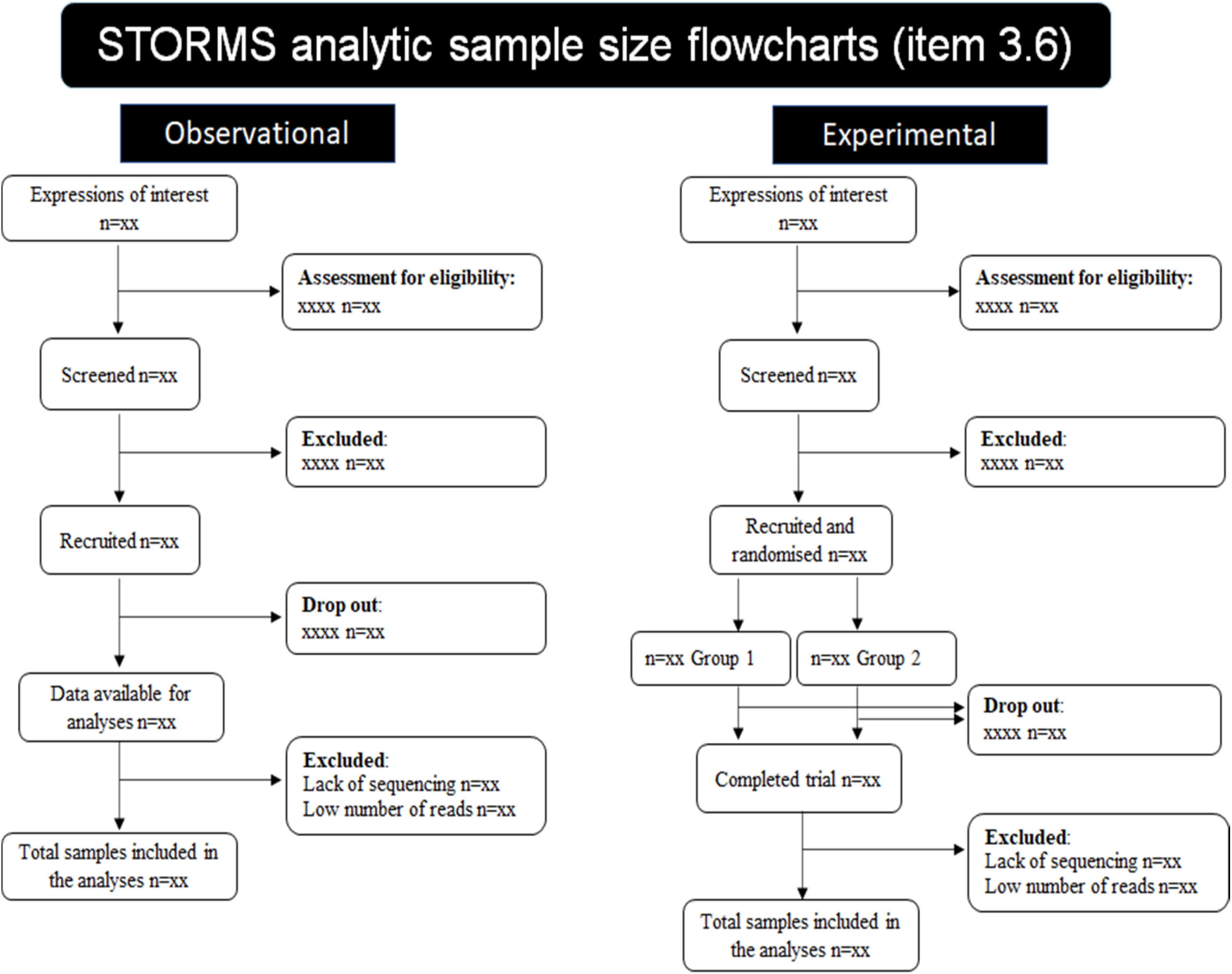
Flowchart templates for item 3.6 (Analytic sample size).

##### Laboratory methods (4.0-4.17)

Since STROBE does not cover laboratory methods, new items were developed for STORMS. Describe laboratory methods in sufficient detail to allow replication. The handling of lab samples should be described, including procedures for sample collection (items 4.1), shipping (item 4.2), and storage (item 4.3).

Because DNA extraction can be a major source of technical differences across studies,(10) DNA extraction methods should be described (item 4.4). Human DNA removal and microbial DNA enrichment, if performed, should also be included (item 4.5). Likewise, if positive controls (item 4.7), negative controls (item 4.8), or contaminant mitigation methods (item 4.9) were used, they should be identified and described.

Sequencing-related methods should be reported. This includes primer selection and DNA amplification (including the variable region of the 16S rRNA gene if applicable) (item 4.6). Major division of sequencing strategy, such as shotgun or amplicon sequencing, should be identified (item 4.11). Finally, the methods used to determine relative abundances should be explained (item 4.12) and the read numbers that serve as denominators recorded.

Batch effects should be discussed as a potential source of confounding, including steps taken to ensure batch effects do not overlap with exposures or outcomes of interest (item 4.13).(29) If conducting metatranscriptomics, metaproteomics, or metabolomics, provide details of those methods (items 4.14 to 4.16).

##### Data sources/measurement (5.0)

For non-microbiome data (e.g. health outcomes, participant socioeconomic, behavioral, dietary, biomedical characteristics, including disease location and activity, and environmental variables), the measurement and definition of each variable should be described (item 5.0). For instance, participant sex and age could be obtained from electronic medical records or from a questionnaire distributed to participants; this data source should be described. Limitations of measurement may also be discussed including potential bias due to misclassification or missing data, as well as any attempts made to address these measurement issues.

##### Research design considerations for causal inference (6.0-6.1)

Observational data is often used to test associations that aim towards causal inference. Methods include, for example, the use of multivariable analysis or matching to adjust for confounding variables between a hypothesized exposure (such as abundance of a microbial taxon) and the disease or condition under study.(30) If variables are adjusted for in the analysis, theoretical justification for inclusion of these variables should be provided (item 6.0). As part of this theoretical justification, consider including a directed acyclic graph showing the hypothesized causal relationships of interest.(31,32) In addition to considering the theoretical motivations for the present study, discuss the potential for selection or survival bias which can distort the observed relationship between the microbiome and variable of interest (item 6.1). For example, such bias may occur due to loss-to-follow-up (in longitudinal studies) or due to participants not being included in the study due to the condition itself (e.g. participants who have died of aggressive forms of colorectal cancer have not survived to be in a hypothetical study of colorectal cancer microbiomes).(33)

##### Bioinformatics and Statistical Methods (7.0-7.9)

Adequate description of bioinformatic and statistical methods is essential to producing a rigorous and reproducible research report. Data transformations (such as normalization, rarefaction, and percentages) should be described (item 7.0). Quality control methods should be fully disclosed so that readers understand criteria for filtering or removing reads or samples (item 7.1). All statistical methods used to analyze the data should be stated, (item 7.3) including how results of interest were selected (e.g. using a p-value, q-value, or other threshold) (item 7.8). Taxonomic, functional profiling, or other sequence analysis methods should be described in detail (item 7.2) In the interest of reproducibility, all software, packages, databases, and libraries used for the pre-processing and analysis of the data should be described and cited including version numbers (item 7.9).

##### Reproducible Research (8.0-8.5)

Reproducible research practices serve as quality checks in the process of publication and further transparency and knowledge sharing, as detailed in the rubric proposed by Schloss.(34) Journals are increasingly implementing reproducible research standards that include the publishing of data and code, and those guidelines should be followed when possible.(35,36) STORMS itemizes the accessibility of data, methods, and code (items 8.0 through 8.5). If possible, data should be deposited into a public repository (items 8.1 and 8.2). If data or code are not or cannot be made publicly available, a description of how interested readers can access the data should be provided. As stated in item 8.0, any protected information should be described along with how such data can be accessed.

#### Results (9.0-10.4)

##### Descriptive Data (9.0)

Descriptive statistics about the study population should be reported (item 9.0). At a minimum, age and sex of all participants should be reported, but other important participant characteristics should be reported when possible including medication use or lifestyle factors such as diet. Consider reporting these data in a descriptive statistics table. Packages such as the table1 package in R make creating such a table straightforward.(37)

##### Outcome Data (10.0-10.4)

The main outcomes of the study should be detailed including descriptive information, findings of interest, and the results of any additional analyses. Descriptive microbiome analysis (for instance, dimension reduction such as Principal Coordinates Analysis, measures of diversity, gross taxonomic composition) should be reported for each group and each time point (item 10.0). This contextualizes the results of differential abundance analysis for readers. When reporting differential abundance test results, the magnitude and direction of differential abundance should be clearly stated (item 10.2) for each identifiable standardized taxonomic unit (item 10.1). Results from other types of analyses such as metabolic function, functional potential, MAG assembly, and RNAseq should be described in the results as well (items 10.3 and 10.4). Additional results (e.g. non-significant results or full differential abundance results) can be included in supplements, but should not be excluded entirely. Although the problem has been known for decades,(35) journals across many fields are recognizing the issue of publication bias and therefore the issue of non-reporting of null results.(38) Including such results in publications will help to reduce the severity of this bias and improve future systematic reviews and meta-analyses.

#### Discussion (11.0-14.0)

Most recommendations for the Discussion section are similar to STROBE including a discussion of the limitations of the present study and associated methods (item 13.0). One additional recommendation is made: discuss the potential for biases and how they would influence the study findings (item 13.1). Many forms of bias such as residual/unmeasured confounding, bias related to compositional analysis,(39) measurement bias, or selection bias(40) could affect the interpretation of the results of the study and it is important to acknowledge potential sources of bias when discussing the results.(41) As described in STROBE, authors should also consider the generalizability of their findings and if these findings could be applicable to the target population or other populations (item 13.2). If different forms of bias were not assessed or assumed to be negligible, this should be stated. Finally, authors should discuss potential future or ongoing research based on findings of the present study (item 14.0).

#### Other Information (15.0-17.0)

In addition to a statement of funding (item 15.0), authors should also include acknowledgements and conflicts of interest statements (items 15.1 and 15.2, respectively). Conflicts of interest statements should be written according to the criteria established by the journal. Finally, the paper should state where supplementary materials and data can be accessed (items 16.0 and 17.0).

## Discussion

The STORMS checklist for reporting on human microbiome studies was developed with the following priorities. The checklist should 1) be easy to understand and use by researchers from various fields, through straightforward use of language and pruning of items rarely relevant to the current literature, 2) be organized in the outline of a manuscript, so it can serve as a tool for authors and for peer reviewers, particularly when included in manuscript submission as a supplemental table with comments, 3) assist in the complete and organized reporting of a study, not in enforcing any particular methods, 4) reuse or modify items from related checklists where relevant, and 5) represent consensus across a broad cross-section of the human microbiome research community. The checklist facilitates manuscript authors in providing a complete, concise, and organized description of their study and its findings. Included as a supplemental table to a manuscript, it also supports efficient peer review and post-publication interpretation.

While other efforts for extending STROBE for microbiome and metagenomic have been proposed (9) and laboratory-focused reporting checklists have been released,(4,42) to our knowledge STORMS is the first comprehensive reporting checklist for human microbiome research. We aim for the STORMS guidelines to improve the quality and transparency of microbiome epidemiology studies by introducing a shared grammar of study reporting in a structured checklist format. Reporting checklists introduced in other disciplines have been shown to improve the quality of journal articles.(5,6)

A major strength of STORMS is the rigor and transparency of its development by a diverse, multidisciplinary consortium of subject matter experts. All feedback received from consortium members, and responses, have been included as Supplement 1. The development of STORMS is an ongoing process, and new versions of the checklist will be released to reflect evolving standards and technological processes. A version control system with changelog has been implemented, and annual reviews of the checklist are planned. Additionally, the working group plans to evaluate STORMS impact on microbiome reporting by examining how many articles are fulfilling checklist items before and after its release. We invite interested readers to join the STORMS Consortium by contacting the corresponding author or by visiting the consortium website for more information (https://www.stormsmicrobiome.org/). We also encourage journals to include the STORMS checklist in their instructions to authors and advise peer reviewers to consult the checklist when reviewing submissions.

There are some limitations to the STORMS checklist. The checklist was not created to assess study or methodological rigor. It is meant to aid authors’ organization and ease the process of reader assessment of how studies are conducted and analyzed. Conclusions about the quality of studies should not be made based on their adherence to STORMS guidelines, although we expect the reporting guidelines to help readers review studies critically. It does not encourage, discourage, or assume the use of null hypothesis significance testing(43) or methods of compositional data analysis(44), topics of some controversy in the field. In general the checklist avoids reference to or guidance on specific statistical methodological decisions.

Through the efforts of the STORMS Consortium working in an iterative, transparent, and collaborative process, the STORMS checklist provides a roadmap for researchers in reporting the results of a human microbiome study. The STORMS Consortium believes that the checklist is sufficiently flexible and user-friendly to support widespread adoption and contribution to microbiome study standards. Its adoption will ideally encourage thoughtful study design, reproducibility, collaboration, and open knowledge sharing between research groups as they explore the human microbiome.

## Notes

http://storms.waldronlab.io/

### Competing Interest Statement

The authors have declared no competing interest.

### Summary of Updates

Updated authorship, added figures, incorporated consortium feedback.

